# Mobility shapes plasmid GC content evolution

**DOI:** 10.64898/2026.07.20.739530

**Authors:** William Matlock, Anna E. Dewar

## Abstract

Plasmids are frequently AT-rich relative to their bacterial hosts. Despite this tendency towards lower GC content, plasmid and host chromosome GC content are positively correlated across diverse collections of plasmid–host pairs. However, the evolutionary processes underlying this pattern remain unclear. The classic model of amelioration predicts that horizontally acquired DNA gradually converges on host nucleotide composition, but since plasmids can repeatedly transfer between bacterial hosts, the opportunity for such host-associated evolution may depend on their transmission dynamics. Using 50,936 plasmid–host pairs from a public sequence database, we found that the apparent global correlation between plasmid and host chromosome GC content was largely driven by differences between bacterial species rather than within species. We therefore accounted for plasmid and host population structure when testing how plasmid mobility shaped host-associated compositional evolution. We compared two contrasting regimes: a population of 3,682 Enterobacterales plasmids distributed across diverse host backgrounds, and six long-term host-associated plasmids from a *Rhizobium leguminosarum* lineage with INSeq-determined gene essentiality data. In the Enterobacterales population, GC content variation was overwhelmingly explained by plasmid lineage rather than host phylogeny, and conjugative plasmids showed greater similarity to their host chromosomes than mobilisable or non-mobilisable plasmids. In the *Rhizobium leguminosarum* plasmids, synonymous-site composition was more similar to the host chromosome among genes required across multiple host life stages. Together, these results support a model in which plasmid mobility influences the opportunity for host-associated evolutionary processes to alter nucleotide composition.

## Introduction

Plasmids are extrachromosomal replicons that are frequently AT-rich relative to their bacterial host chromosomes [1]. Despite this tendency towards lower GC content, plasmid and host chromosome GC content are positively correlated across large collections of plasmid-host pairs [1]. This correlation has traditionally been interpreted as evidence of amelioration, whereby horizontally acquired DNA gradually acquires a nucleotide composition more similar to the host chromosome through shared mutational processes and selective pressures [2]. However, several non-mutually exclusive processes may shape plasmid nucleotide composition. Although a genome-wide mutational bias toward AT substitutions would drive both plasmids and chromosomes towards lower GC content over time [3], differences in inheritance, transmission dynamics, and effective population sizes may cause plasmids to follow distinct evolutionary trajectories even under a shared mutational spectrum [4]. In addition, selection on plasmid maintenance and expression within hosts may further influence nucleotide composition [5, 6, 7, 8]. Therefore, a key question is not simply whether plasmids ameliorate, but under what evolutionary conditions plasmids have the opportunity to undergo host-associated compositional evolution.

Although plasmids depend on host replication and repair machinery, frequent horizontal transfer repeatedly exposes them to different bacterial backgrounds, reducing the opportunity for plasmid nucleotide composition to track any individual host lineage. Conversely, plasmids with prolonged associations with particular hosts may experience greater opportunity for host-associated evolutionary processes. Consistent with this possibility, analyses of trinucleotide signatures in public plasmid sequences suggest that replicon types associated with narrower host ranges tend to more closely resemble host GC composition [9]. However, the relationship between mobility and host association is not straightforward. Recent work suggests that conjugative plasmids can exhibit stronger host-lineage associations than mobilisable plasmids, despite encoding the machinery required for autonomous transfer [10]. This indicates that the evolutionary consequences of mobility depend on realised transmission dynamics and the stability of host associations.

A growing body of evidence suggests that nucleotide composition may represent a stable property of plasmid lineages. GC content is often conserved within plasmid groups [11], and has been associated with increased antimicrobial resistance gene carriage [12] and epidemiological success in human bloodstream infections [13]. Such group-level conservation means that plasmid-host GC correlations may reflect shared ancestry rather than ongoing host-associated evolution. Hence, testing the role of mobility requires accounting for both plasmid and host population structure.

In this study, we first revisited the public sequence database analysis of Almpanis et al. (2018) [1] to determine whether the classic plasmid-host GC content correlation persisted after accounting for population structure. We then investigated whether mobility influenced the extent of host-associated compositional evolution using two contrasting evolutionary systems that differ in the extent of horizontal transfer and long-term host association. First, we examined a diverse population of clinical and non-clinical Enterobacterales isolates from Oxfordshire, previously shown to host plasmid lineages shared across genera and ecological niches [14]. Second, we examined a collection of *Rhizobium leguminosarum* plasmids with a long-term host association [15].

## Materials and methods

### PLSDB analysis

We first downloaded the metadata from PLSDB (v. 2024_05_31_v2), a curated, non-redundant subset of NCBI diversity [16]. We then downloaded the genome assemblies using NCBI datasets (v. 18.30.1; `download genome accession` and `datasets rehydrate`), only retaining assemblies that were complete (`ASSEMBLY_Status` was `Complete Genome`, which required a gapless chromosome and no more than 9 ambiguous bases in a row) and where the plasmid was circularised (`NUCCORE_topology` was `circular`), and calculated the GC content (see `plsdb_model.R`).

### Enterobacterales population analysis

We retrieved a total of 1,454 genome assemblies from Matlock et al. (2023), which contained 3,682 plasmids [14]. Four assemblies from the original study were removed because the chromosomes appeared fragmented; all remaining replicons were fully circularised. Bioinformatic analyses were performed using a custom Nextflow pipeline with Docker containers [17, 18]. Replicon length and GC content were calculated using a custom Python script. Plasmid mobility was predicted using MOB-typer (v. 3.1.9; default parameters) [19]. Plasmid communities and subcommunities were determined using pling (v. 3.0.2; `--batch_size 2000`) [20]. The chromosomal tree was constructed using mashtree (v. 1.4.6; `--mindepth` 0) [21]. Taxonomic assignments were determined using mlst (v. 2.28.1; default parameters) [22, 23].

We modelled plasmid GC content minus host chromosome GC content (ΔGC) using a Bayesian linear phylogenetic mixed-effects model of the form

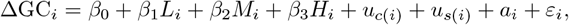

where *L*_*i*_ was the centred log_10_-scaled plasmid length, *M*_*i*_ was the predicted plasmid mobility class (conjugative, mobilisable, or non-mobilisable), and *H*_*i*_ was the centred host chromosome GC content. Random intercepts were included for plasmid community (*u*_*c*(*i*)_) and subcommunity nested within community (*u*_*s*(*i*)_) to account for hierarchical plasmid population structure. Host evolutionary relatedness was modelled using a phylogenetic random effect (*a*_*i*_) with covariance determined by the chromosomal tree. Under this formulation, the coefficient for host chromosome GC content quantified the extent to which plasmid GC content tracked host GC content after accounting for plasmid and host population structure.

If plasmid GC increased by one percentage point for every one percentage point increase in host GC, ΔGC would remain constant and the coefficient would approach zero. Conversely, a coefficient approaching *−*1 would indicate that plasmid GC remained approximately constant across hosts with different chromosome GC contents.

All models were implemented in R using the `brms` package (see `enterobacterales_model.R`) [24, 25]. Models were fit using four chains, each with 2,000 warm-up and 2,000 sampling iterations. Convergence was assessed using 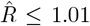, all tail and bulk effective sample sizes ≥ 200, and inspection of trace plots. We excluded subcommunities with fewer than five members because variance estimates for very small groups are poorly informed and may be dominated by sampling noise. Continuous predictors (log_10_-scaled plasmid length and host chromosome GC content) were mean-centred to improve Markov chain Monte Carlo (MCMC) sampling efficiency and to ensure that the global and group-level intercepts represent the expected ΔGC at the average values of the sample. These continuous predictors were initially modelled with smoothing functions to assess non-linearity. However, conditional effect plots indicated that the posterior estimates for these continuous predictors had 95% credible intervals consistent with approximately linear relationships (Figure S1). Therefore, smooth terms were replaced with linear terms in the final models to avoid overfitting.

To quantify the proportion of total variance in ΔGC explained by each hierarchical clustering level and host phylogeny, we calculated Bayesian intraclass correlation coefficients (ICCs). The total variance 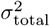 was decomposed as

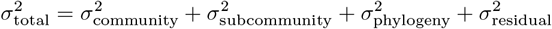

where 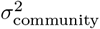 was the variance across plasmid communities, 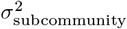 was the variance across sub-communities nested within communities, 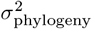 was the variance associated with host phylogenetic relationships (modelled via the host chromosome covariance matrix), and 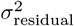 was the residual (unexplained) variance. The ICC for any given variance component *k* ∈ {community, subcommunity, phylogeny} was defined as

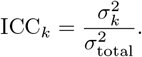

We calculated these ratios at each iteration of the MCMC chains, allowing us to propagate uncertainty and derive full posterior distributions and hence 95% credible intervals.

### *Rhizobium leguminosarum* analysis

Wheatley, R. M. et al. (2020) used mariner-based transposon insertion sequencing (INSeq) to determine plasmid gene essentiality within *R. leguminosarum bv. viciae* Rlv3841 (also called *R. johnstonii* 3841) across four stages of the symbiotic lifestyle (free-living growth in the rhizosphere, root attachment, differentiation into bacteroids, and release from nodules) [15]. We recovered essentiality data for 2,344 genes across the six plasmids. The reference genome annotation (GCF 000009265.1) was downloaded in GBFF format using the NCBI datasets tool (v. 15.27.1). We developed the custom script `rhizobium_gc.py` to calculate the GC content at the first, second and third codon positions (GC1–3) together with four-fold degenerate sites (GC4) for every annotated gene.

For each gene, we calculated ΔGC1-4 by subtracting host chromosome GC content (61.09) from GC1, GC2, GC3 and GC4. Then, for each gene *i*, we jointly modelled the four response variables using a Bayesian multivariate linear mixed-effects model

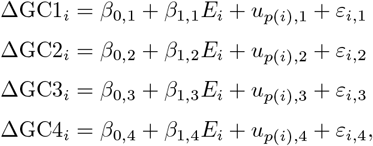

where *E*_*i*_ was the gene essentiality score (0–4), and *u*_*p*(*i*)_ represented a plasmid-level random intercept accounting for shared evolutionary history among genes located on the same plasmid. The residual terms were modelled jointly to allow for correlations between codon-position-specific deviations after accounting for gene essentiality and plasmid identity.

The model was implemented in R using the `brms` package (see `rhizobium_model.R`) [24, 25]. Models were fit using four chains, each with 2,000 warm-up and 2,000 sampling iterations. Convergence was assessed using 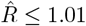, all tail and bulk effective sample sizes *≥* 200, and inspection of trace plots.

### Data availability

Data needed to reproduce the PLSDB analysis are stably archived on Figshare under the DOI https://doi.org/10.6084/m9.figshare.27252609.v2. Data needed to reproduce the Enterobacterales analysis (genome assemblies and sample sheet) and *R. leguminosarum* analysis (NCBI sequence data and original study metadata) will be stably archived on Zenodo. There, we will also archive the outputs used for figures and modelling (PLSDB plasmid-host GC content values, all results from the Enterobacterales Nextflow pipeline, and *R. leguminosarum* GC content values).

### Code availability

The results from this study can be reproduced by following the instructions in https://github.com/wtmatlock/plasmid-gc, which will be stably archived on Zenodo.

## Results

### Plasmid-host GC content correlation is confounded by population structure

Almpanis et al. (2018) analysed 5,744 non-redundant plasmid-host pairs from NCBI, finding a strong Pearson correlation coefficient of 0.74 between plasmid and host chromosome GC content [1]. We replicated this statistical test using 50,936 non-redundant plasmid-host pairs from PLSDB (representing a total of 20,349 genome assemblies), finding a Pearson correlation coefficient of 0.917 (95% confidence interval: 0.916 to 0.918; *t*(50,934) = 518.53; *p*-value < 0.0001). However, inspection of the raw data (Figure 1) suggested that this global correlation was largely driven by variation between species. Among densely sampled species, plasmid GC content showed little evidence of tracking with host chromosome GC content. For example, restricting the analysis to the most represented species, *E. coli* (10,039/50,936 plasmid-host pairs), the correlation remained statistically significant but was negligible in magnitude (0.046; 95% confidence interval: 0.026 to 0.065; *t*(10,037) = 4.567; *p*-value < 0.0001). This motivated a statistical approach that accounted for population structure in both hosts and plasmids [26]. Moreover, public sequence repositories aggregate genomes from disparate ecological contexts, often biased by specific pathogens and outbreaks. We therefore sought to test these patterns within an epidemiologically coherent population of plasmids and hosts.

**Figure 1:**
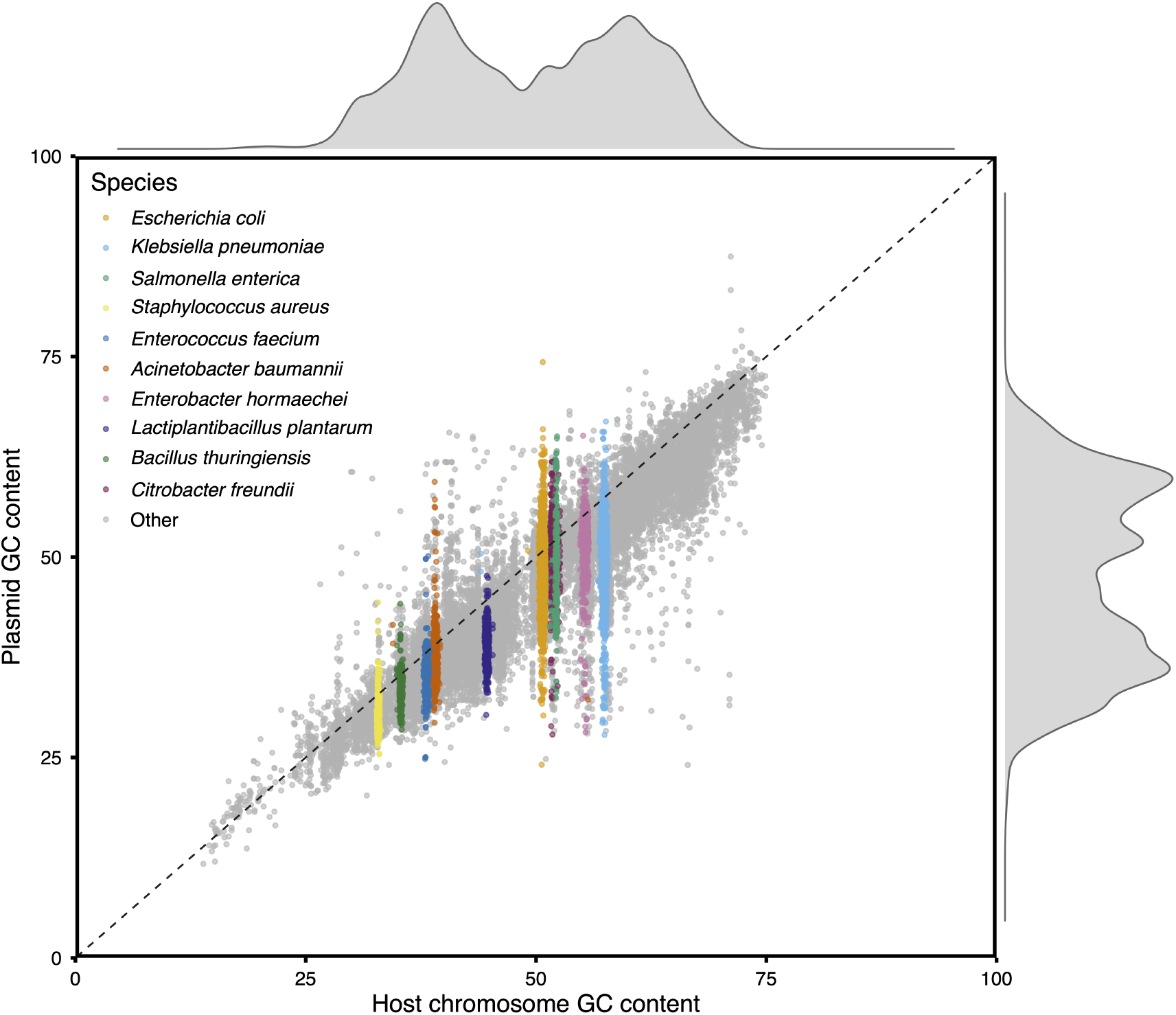
Comparison of plasmid and host chromosome GC content across publicly available sequence diversity. The dataset comprised 50,936 non-redundant plasmid–host pairs. Host species are highlighted for the ten most frequently represented taxa; all remaining species are shown in grey. The dashed line represents equality between plasmid and host chromosome GC content.

### Enterobacterales plasmid GC content is structured by plasmid lineage, not host ancestry

Enterobacterales plasmids are highly diverse and often mobile [27, 14]. We reused a dataset of 3,682 plasmids originating from 1,454 isolates, representing seven Enterobacterales genera from at least nine species (Table 1). Isolates were sampled from human bloodstream infections, livestock, and wastewater, all within 60km in Oxfordshire between 2008 and 2020. This provided a geographically and temporally coherent snapshot of an Enterobacterales plasmid population.

**Table 1:**
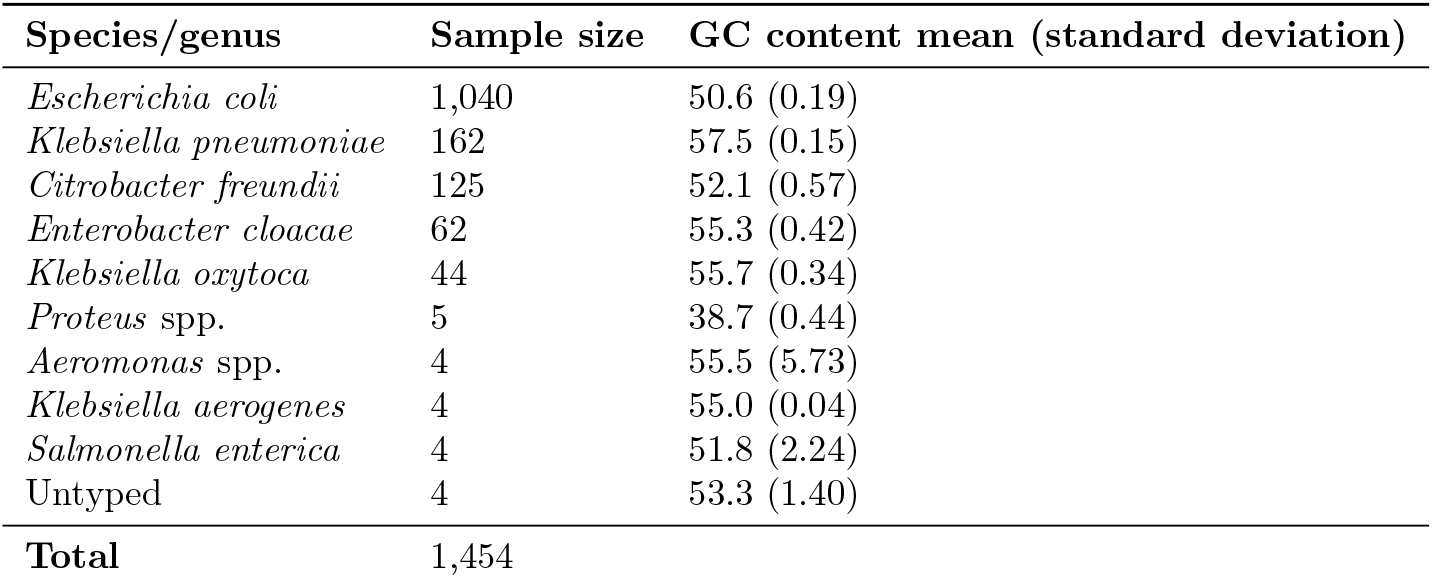
Enterobacterales dataset summary by species or genera (typed by mlst; see Methods).

| Species/genus | Sample size | GC content mean (standard deviation) |
| --- | --- | --- |
| *Escherichia coli* | 1,040 | 50.6 (0.19) |
| *Klebsiella pneumoniae* | 162 | 57.5 (0.15) |
| *Citrobacter freundii* | 125 | 52.1 (0.57) |
| *Enterobacter cloacae* | 62 | 55.3 (0.42) |
| *Klebsiella oxytoca* | 44 | 55.7 (0.34) |
| *Proteus* spp. | 5 | 38.7 (0.44) |
| *Aeromonas* spp. | 4 | 55.5 (5.73) |
| *Klebsiella aerogenes* | 4 | 55.0 (0.04) |
| *Salmonella enterica* | 4 | 51.8 (2.24) |
| Untyped | 4 | 53.3 (1.40) |
| **Total** | 1,454 |  |

To characterise the plasmid population structure, we resolved the 3,682 plasmids into genetically related communities and subcommunities. These groupings were based on structural similarity, specifically the sharing and arrangement of homologous sequence blocks, rather than overall nucleotide identity. We excluded subcommunities containing fewer than five plasmids, leaving 2,305 plasmids across 89 subcommunities for analysis (Figure 2a). For each plasmid-host pair, we then calculated plasmid GC content minus host chromosome GC content (ΔGC; Figure 2b). Negative values indicated that plasmids were AT-rich relative to their host chromosomes. We then predicted ΔGC using plasmid length, predicted mobility, and host chromosome GC content, accounting for plasmid and host population structure.

**Figure 2:**
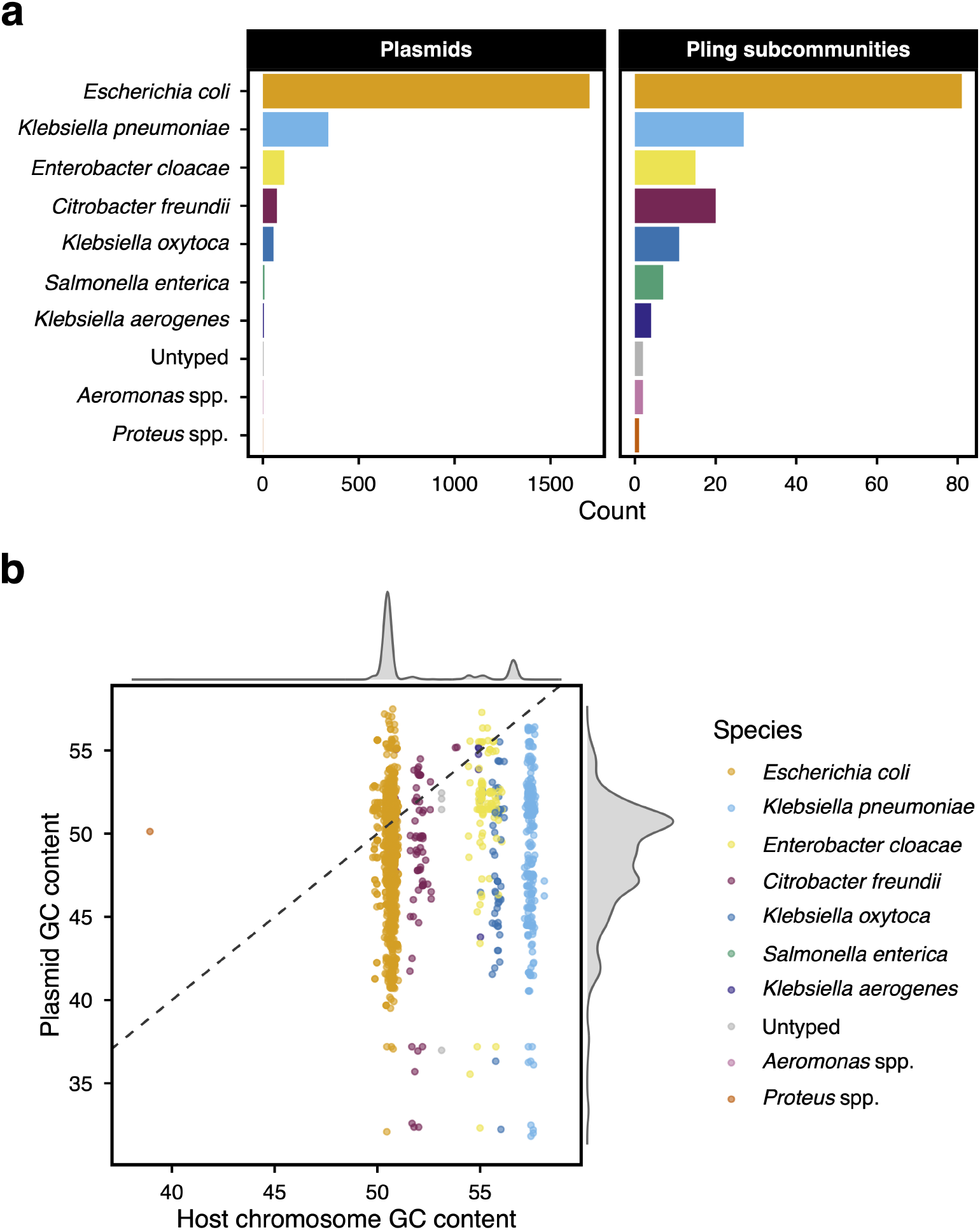
A natural population of Enterobacterales plasmids. Here we present the 2,305/3,682 plasmid-host pairs used in the model, originating from pling subcommunities with at least 5 members. The number of plasmids and pling subcommunities stratified by host species. **(b)** The GC content of the plasmids and their host chromosomes. The dashed line represents equality between host chromosome and plasmid GC content.

While plasmids were, on average, 4.10% more AT-rich than their hosts (95% credible interval (CI): –6.79 to –1.36), ΔGC was largely structured by plasmid-intrinsic factors. A tenfold increase in plasmid length was associated with a 1.03% increase in ΔGC (95% CI: 0.64 to 1.40). Compared with conjugative plasmids, both mobilisable and non-mobilisable plasmids were progressively more AT-rich (–1.01, 95% CI: –1.55 to –0.47; and –2.18, 95% CI: –2.69 to –1.66, respectively). Thus, conjugative plasmids showed the greatest similarity to host chromosome GC content.

The estimated effect of host chromosome GC content was close to –1 (–1.03, 95% CI: –1.27 to –0.84), indicating that, after accounting for population structure, plasmid GC content was largely independent of host GC background. Consistent with this, Bayesian intraclass correlation coefficients (ICCs) showed that plasmid community and subcommunity together explained 93.1% (95% CI: 88.4 to 96.4) of the total variance in ΔGC, whereas host phylogeny explained only 0.63% (95% CI: 0.16 to 1.62). Together, these results indicated that plasmid GC content was more strongly associated with plasmid lineage than host ancestry.

It was possible that the strong community- and subcommunity-level variance simply reflected low within-group diversity, rather than meaningful structure. This concern was greatest for very small plasmid groups, which can be highly conserved [14, 28]. To address this, we re-ran the model on the 27/89 subcommunities (1,530/2,073 plasmids) with a GC content standard deviation of at least 1. This threshold far exceeded the variation in chromosomal GC content observed for the densely sampled species (Table 1). The 95% CIs for the ICCs remained consistent with the full model, as community and subcommunity accounted for 84.87% (95% CI: 68.89 to 95.30) of the variance in ΔGC, whereas host phylogeny accounted for 0.90% (95% CI: 0.03 to 3.75).

### *Rhizobium leguminosarum* plasmid synonymous-site composition varies with gene essentiality

If frequent plasmid transmission limits host-associated compositional evolution, then long-term host association should facilitate it. *R. leguminosarum bv. viciae* carries six plasmids within a long-term host-associated symbiotic lineage, providing a contrasting evolutionary system in which plasmids experience a relatively stable host environment. We further hypothesised that genes required across more lifestyle stages may show stronger host-associated codon adaptation, since selection acting on translational efficiency may favour compatibility with host tRNA pools [29, 30].

To test these hypotheses, we used gene essentiality classifications for 2,344 genes across the six plasmids, determined by mariner-based transposon insertion sequencing (INSeq) across four stages of the symbiotic lifestyle (free-living growth in the rhizosphere, root attachment, differentiation into bacteroids, and release from nodules) [15]. Each gene was assigned an essentiality score from 0 to 4, corresponding to the number of lifestyle stages in which it was required. We calculated relative gene-wise GC content at the first, second, and third codon positions, as well as four-fold degenerate sites, by subtracting the host chromosome GC content from each measure (ΔGC1–4; Figure 3). These response variables were jointly modelled as a function of gene essentiality score (0–4), while accounting for variation between plasmids. Because synonymous sites experience weaker protein-level constraint than nonsynonymous sites, we expected the strongest signatures of host-associated compositional evolution at GC3 and GC4 [31].

**Figure 3:**
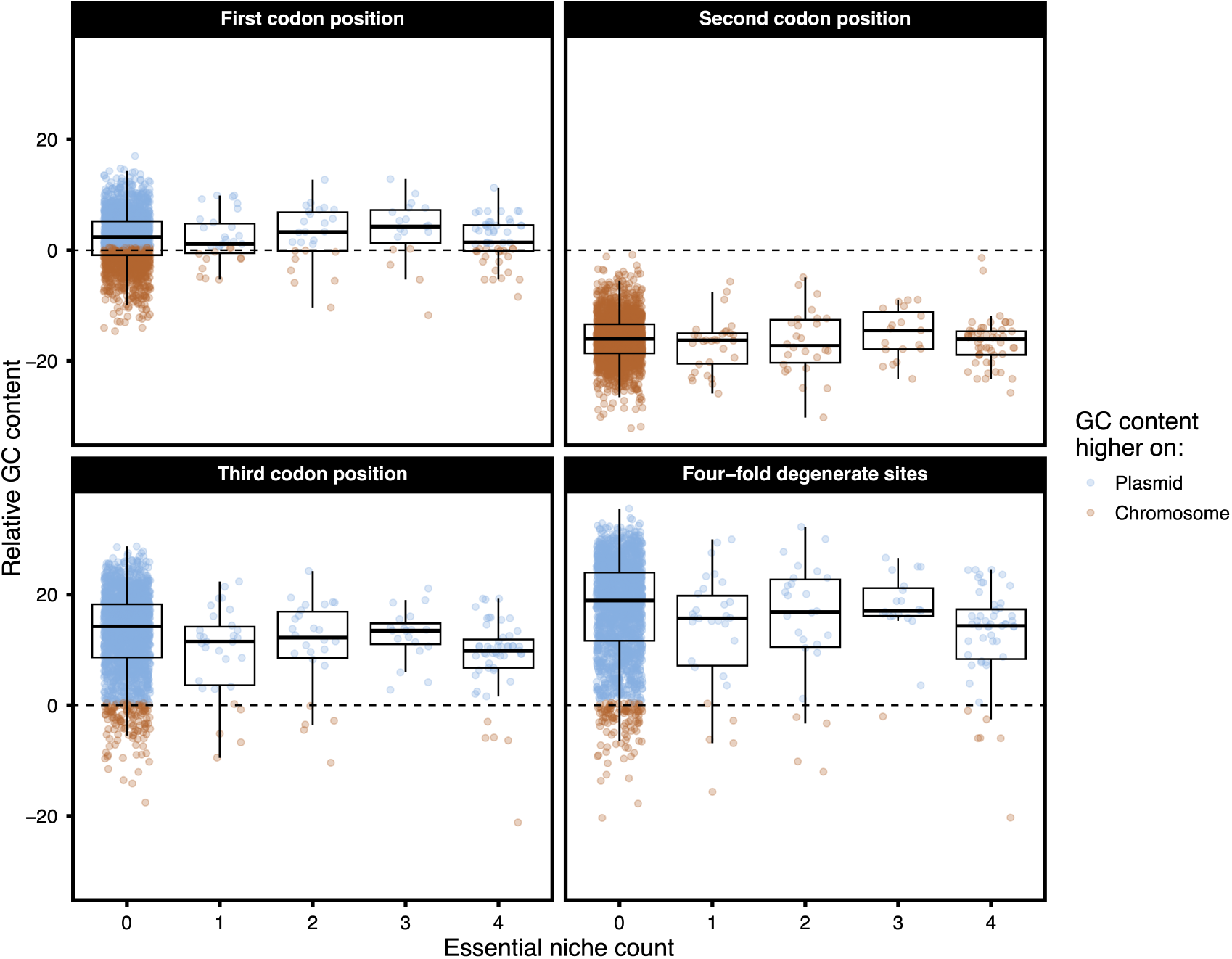
Plasmid GC content bias at each codon position in *Rhizobium leguminosarum*, stratified by gene essentiality. Relative GC content was defined as the difference between plasmid gene-level GC content and the host chromosome GC content (61.09), calculated separately for first (GC1), second (GC2), and third (GC3) codon positions, as well as four-fold degenerate sites (GC4). Gene essentiality reflects the number of lifestyle stages (0–4) in which a gene was required, as determined experimentally using mariner-based transposon insertion sequencing (INSeq).

For the more constrained first and second codon positions (ΔGC1 and ΔGC2), genes with an essentiality score of zero were 1.88% GC-rich (95% CI: 1.17 to 2.43) and 15.99% AT-rich (95% CI: 15.73 to 16.23), respectively, relative to the host chromosome. Neither position showed a clear association with essentiality score (ΔGC1 slope: 0.12, 95% CI: –0.16 to 0.41; ΔGC2 slope: –0.13, 95% CI: –0.39 to 0.12). In contrast, for the less constrained ΔGC3, genes with an essentiality score of zero were 11.64% GC-rich (95% CI: 8.13 to 15.19), but this value decreased with increasing essentiality score (–0.44, 95% CI: –0.86 to –0.02). We also detected a strong positive residual correlation between ΔGC3 and the least constrained ΔGC4 (0.93, 95% CI: 0.92 to 0.93). Together, these results were consistent with host-associated compositional evolution acting primarily at synonymous sites.

We also detected substantial heterogeneity between plasmids in their baseline synonymous-site GC composition. The estimated among-plasmid standard deviation was 0.60% (95% CI: 0.08 to 1.51) for ΔGC1 and 0.15% (95% CI: 0.00 to 0.53) for ΔGC2, but increased to 4.15% (95% CI: 2.13 to 8.25) for ΔGC3 and 5.43% (95% CI: 2.83 to 10.67) for ΔGC4. Thus, despite sharing a common host background, plasmids differed substantially in their synonymous-site composition, suggesting that the extent of host-associated compositional evolution varied between plasmid lineages.

## Discussion

In this study, we argue that plasmid mobility influences the opportunity for host-associated compositional evolution. The classic model of amelioration posits that horizontally acquired DNA may gradually acquire nucleotide signatures similar to those of its host due to shared mutational biases and selective pressures [2]. However, our results show that this is not a universal outcome for plasmids; rather, the opportunity for amelioration-like processes is constrained by their transmission dynamics. In the Enterobacterales population, plasmid lineages distributed across diverse bacterial backgrounds showed GC content patterns overwhelmingly structured by plasmid lineage, providing limited evidence for host-associated compositional tracking [14]. However, conjugative plasmids showed greater similarity to their host chromosomes than mobilisable plasmids. Although initially counterintuitive, this result highlights that transfer potential does not necessarily translate into promiscuity. Mobilisable plasmids may exploit diverse conjugative elements, whereas some conjugative plasmid lineages may remain restricted to particular host backgrounds [10].

Conversely, in the more stable *R. leguminosarum* system, synonymous-site composition varied with gene essentiality, with genes required across multiple life stages showing greater similarity to host chromosome composition. This signal also varied between plasmids, suggesting that the extent of host-associated compositional change differs between plasmid lineages. This is consistent with recent population genomic analyses showing contrasting transmission regimes among *R. leguminosarum* plasmids, including those carrying key symbiosis genes [32].

The strong global correlation observed in the PLSDB analysis represented a Simpson’s paradox-like pattern, where an association observed after pooling groups was driven primarily by differences between groups rather than consistent relationships within groups. Here, species-level differences in chromosome GC content generated a strong overall plasmid-host correlation, despite limited GC content tracking within species. This highlights the importance of accounting for population structure in such models.

This study has several limitations. First, GC content provides an indirect measure of long-term evolutionary processes and cannot by itself distinguish among the contributions of mutation bias, drift, and selection. Therefore, our results should be interpreted as evidence that mobility influences the extent to which host-associated processes shape plasmid nucleotide composition. Second, plasmid GC content reflects both nucleotide-level evolutionary processes and differences in gene content acquired through horizontal transfer. Although accounting for plasmid population structure reduces the likelihood that these patterns simply reflect broad lineage differences, we cannot exclude contributions from lineage-specific accessory gene repertoires. Third, plasmid mobility was inferred *in silico*, which, whilst often accurate, is inferior to experimental phenotyping. Fourth, we used gene essentiality as a proxy for functional constraint. Whilst genes with higher essentiality scores are required across more lifestyle stages, they are not necessarily universally essential or subject to identical selective pressures. Lastly, since we analysed extant sequence composition rather than evolutionary trajectories, our analysis cannot distinguish active compositional change from historical phylogenetic inertia.

Overall, this study supports mobility as an important factor shaping plasmid compositional evolution. Plasmids that repeatedly transfer among diverse hosts retain lineage-specific nucleotide signatures, whereas long-term host association provides the opportunity for host-associated processes to reshape nucleotide composition. More broadly, these findings suggest that understanding the evolution of mobile genetic elements requires considering their transmission dynamics and the stability of their host associations.

## Funding information

WM was supported by the Wellcome Early-Career Award 319534/Z/24/Z. AD was supported by a Career Development Research Fellowship from St. John’s College, University of Oxford.

## Acknowledgements

This study arose from discussions at the EMBO Workshop on ‘Plasmids as Vehicles of AMR Spread’, 18–23 September 2023. We are grateful to the organisers and attendees. We also thank Edward Feil and Craig MacLean for the valuable discussions.

## Conflicts of interest

The authors have no conflicts of interest.

**Figure S1:**
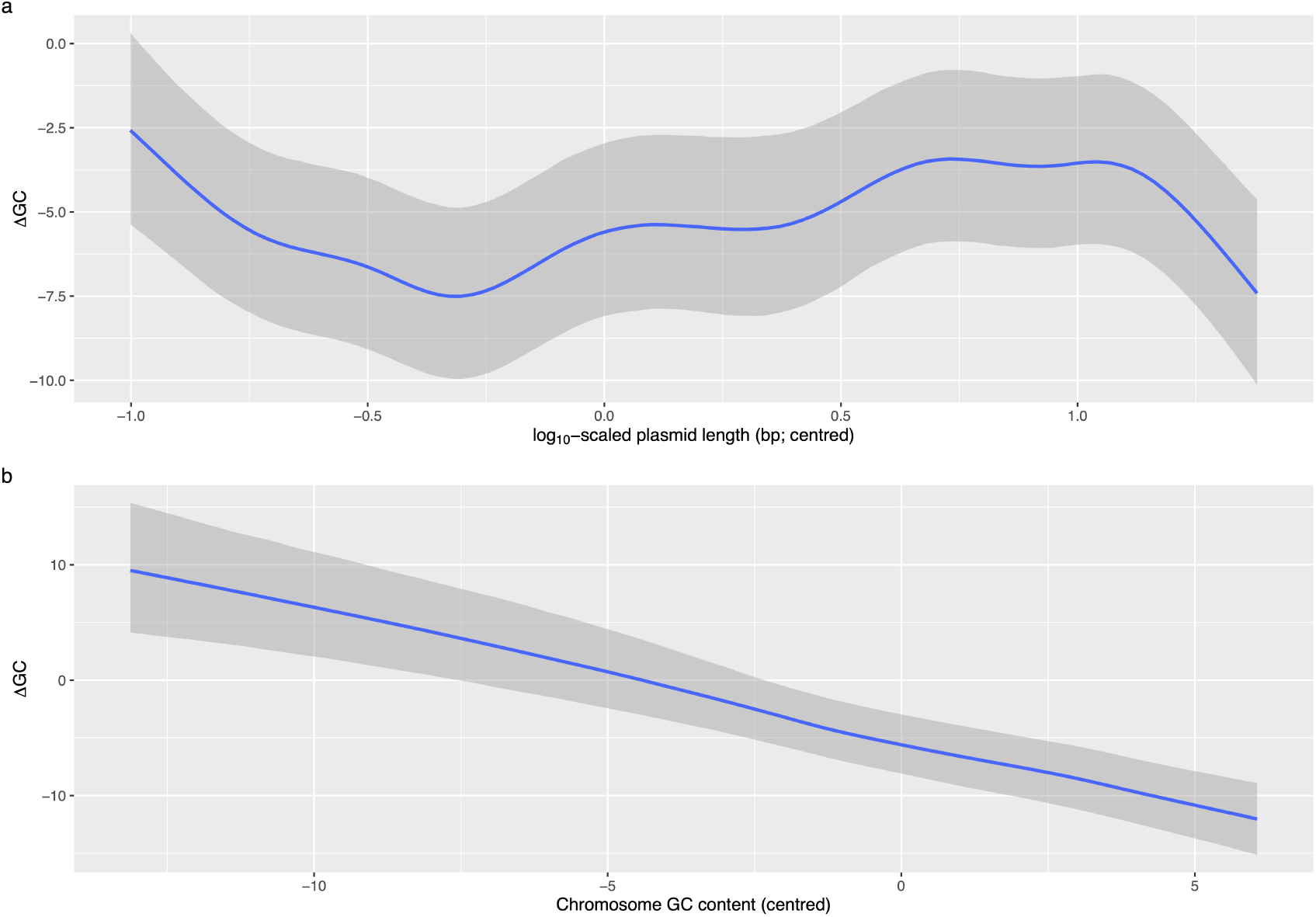
Conditional smooth effects of plasmid length and host chromosome GC content on ΔGC (plasmid GC – host chromosome GC). Conditional effects are plotted from the Enterobacterales model, showing the partial relationship of each predictor while holding all other covariates constant at their reference or mean values. **(a)** The conditional effect of centred, log_10_-scaled plasmid length (bp) on ΔGC. **(b)** The conditional effect of centred host chromosome GC content on ΔGC. In both panels, the solid line represents the posterior mean estimate, and the shaded area denotes the 95% credible interval (CI). In both cases, the 95% CIs are consistent with linearity.

## Notes

https://github.com/wtmatlock/plasmid-gc

### Competing Interest Statement

The authors have declared no competing interest.

### Summary of Updates

We have revised the Introduction and corrected a few typos throughout.

